# Somatic mutational profiles and germline polygenic risk scores in human cancer

**DOI:** 10.1101/2021.01.28.428663

**Authors:** Yuxi Liu, Alexander Gusev, Yujing J. Heng, Ludmil B. Alexandrov, Peter Kraft

## Abstract

The mutational profile of a cancer reflects the activity of the mutagenic processes which have been operative throughout the lineage of the cancer cell. These processes leave characteristic profiles of somatic mutations called mutational signatures. Mutational signatures, including single-based substitution (SBS) signatures, may reflect the effects of exogenous or endogenous exposures. Here, we used polygenic risk score (PRS) as proxies for exposures and examined the association between somatic mutational profiles and germline PRS in 12 cancer types from The Cancer Genome Atlas project. We found 17 statistically significant associations after Bonferroni correction (p < 3.15×10^−5^), including positive associations between germline inflammatory bowel disease PRS and number of somatic mutations of signature SBS1 in prostate cancer and APOBEC-related signatures in breast cancer. The age at menarche PRS was inversely associated with mutation counts of SBS1 in prostate cancer. Our analysis suggests that there are robust associations between tumor somatic mutational profiles and germline PRS. These may reflect mechanisms through hormone regulation and immunological responses that contribute to cancer etiology and drive cancer progression.

## INTRODUCTION

Cancer is driven by the accumulation of somatic mutations. In contrast to germline variants, which are inherited from egg or sperm and occur in DNA of every cell in the body, somatic mutations are generated from mutational processes of exogenous and endogenous exposures as well as DNA enzymatic modifications and failure/infidelity of DNA repair and replication^1, 2, 3^. Mutational processes result in different mutation types (e.g., C>T substitution at the mutated base of ACG motif) with characteristic combinations of mutation types constituting different mutational signatures^1, 4^. Previous studies have identified and confirmed more than 50 distinct signatures of single-base substitution (SBS) derived from the analysis of whole genome and whole exome sequences (WES) of multiple cancer types^1, 4, 5, 6, 7, 8, 9, 10, 11, 12, 13, 14, 15, 16^, but the etiologies for many of these signatures remain largely unexplored. In addition, tumor mutational burden (TMB), which quantifies the total mutations per megabase in a tumor tissue, has been suggested as a biomarker to predict the response of a patient to immunotherapy^17, 18, 19^. Recently, The US Food and Drug Administration approved the use of pembrolizumab, a humanized antibody for cancer immunotherapy, in patients with TMB-High solid tumors. However, currently, it is not fully understood the reason most patients with high TMB benefit from immunotherapy.

Mutational signatures reflect the activity of the mutational processes that have been active through a person’s life^2, 20, 21, 22^. The identified mutational signatures reflect both processes commonly found across cancer types as well as processes confined to a particular cancer type. For example, signature SBS2 and SBS13, both attributed to the enzymatic activity of the APOBEC family of cytidine deaminases, are present in multiple cancer types^6^. In contrast, signature SBS12, whose etiology is still unknown, is almost exclusively found in liver cancers^1^. Some signatures reflect the effects of lifestyle choices, such as signature SBS4, which is associated with tobacco smoking in multiple cancer types, or environmental exposures, such as signatures SBS7a/b/c/d which are imprinted by exposure to ultraviolet light^1, 23^. Some signatures are caused by endogenous exposures, for example, the clock-like signature SBS1 is attributed to endogenous deamination of 5-methylcytosine^24^. However, the etiologies for many other recently identified signatures remain unclear. Linking those signatures of unknown origin to cancer risk factors may suggest mechanism and provide avenues for further investigation.

Previous studies of multiple cancer types have found associations between somatic mutational burden and germline genetic risk factors^25^. For example, germline *MC1R* R alleles carrier status is significantly associated with somatic mutational burden in melanomas^26^. Germline statuses of *ZNF750* and *CDC27* have an impact on somatic mutational signatures in esophageal squamous cell carcinomas^27^. rs2588809 carrier status at gene *RAD51B* is significantly associated with TMB in breast cancer^28^. rs17000526, a variant associated with *APOBEC3B* expression, is strongly associated with APOBEC signature mutations in bladder cancer^29^. Carter et al.^30^ investigated the interaction between an array of germline variants and somatic events across cancers and found robust associations. In addition to studying the association at the level of individual variants, one study also examined the relationship between polygenic risk score (PRS), which combines the effect of multiple germline variants, and somatic mutational burden. They found that germline PRS of breast cancer was inversely associated with total somatic mutation counts in breast tumor samples^28^, but the underlying mechanism is still obscure.

Here, we performed a pan-cancer analysis of the association between tumor somatic mutational profiles and germline PRS of cancers and non-cancer traits using data from The Cancer Genome Atlas (TCGA). Studies with comprehensive somatic mutation data do not always have complete and accurate epidemiological exposure data. PRS can be used as instruments for those unmeasured exogenous and endogenous risk factors. Studying the relationship between germline PRS and somatic mutations can also provide insight into the underlying biological mechanisms of cancer development.

## RESULTS

### TCGA germline and somatic data

We obtained tumor somatic mutation data, germline variant data, and clinical data for 5,766 patients of European ancestry across 12 cancer types from TCGA. The sample size for each cancer type ranged from 333 (kidney renal clear cell carcinoma) to 929 (breast invasive carcinoma). A mutational signature was included in the analysis if there was at least one cancer type where both more than 20 samples and more than 20% of the samples in the cancer type had mutations attributed to that signature. Overall, somatic data were retrieved for (Fig. 1): *(i)* total somatic mutation counts (TSMC), defined as the total number of somatic missense mutations^31^; *(ii)* nine individual SBS mutational signatures; and *(iii)* one combined APOBEC-related signatures (SBS2 and SBS13).

**Fig 1.**
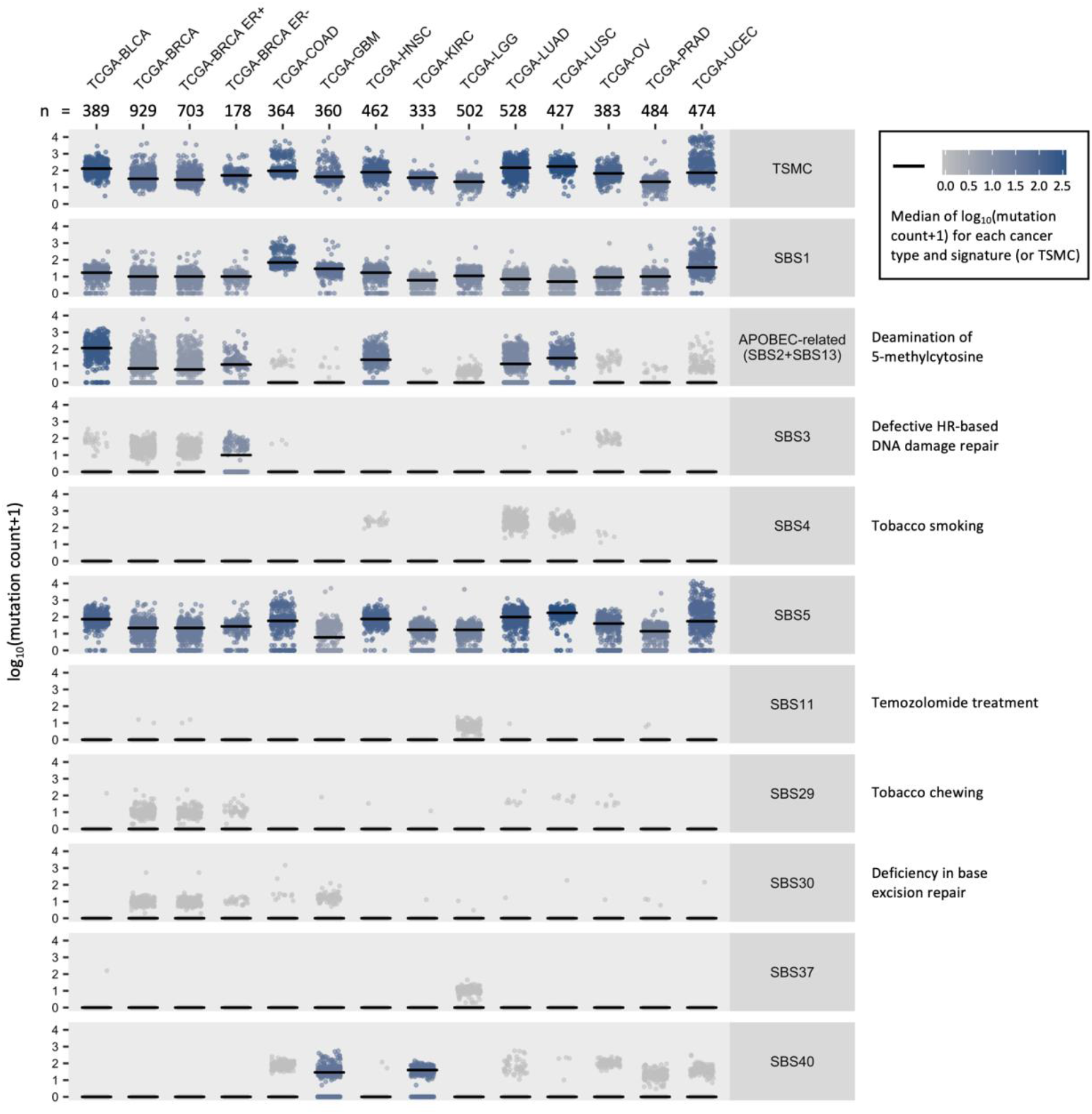
Mutation counts of SBS signatures and TSMC across 12 TCGA cancer types and two subtypes of breast cancer. Each dot represents a tumor sample. The median of log_10_(mutation count+1) for each cancer type and SBS signature (or TSMC) is represented by both the color of points and the black short lines in each panel. Sample size for each cancer type is shown on the top. Abbreviations: TSMC = total somatic mutation counts; SBS = single-base substitution; BLCA = bladder urothelial carcinoma; BRCA = breast invasive carcinoma; BRCA ER+ = breast invasive carcinoma with estrogen receptors; BRCA ER- = breast invasive carcinoma without estrogen receptors; COAD = colon adenocarcinoma; GBM = glioblastoma multiforme; HNSC = head and neck squamous cell carcinoma; KIRC = kidney renal clear cell carcinoma; LGG = brain lower grade glioma; LUAD = lung adenocarcinoma; LUSC = lung squamous cell carcinoma; OV = ovarian serous cystadenocarcinoma; PRAD = prostate adenocarcinoma; UCEC = uterine corpus endometrial carcinoma.

### Calculation of germline PRS

We calculated germline PRS for 23 cancers and non-cancer traits, including the 12 selected cancer types, breast cancer stratified into estrogen receptor positive (ER+) and negative (ER-) subtypes, cancer risk determinants (cigarettes per day, drink per week, and body mass index (BMI)), hormonal factors (age at menarche and age at natural menopause), and immune-mediated inflammatory diseases (inflammatory bowel disease (IBD), ulcerative colitis (UC), Crohn’s disease (CD), and rheumatoid arthritis (RA)) using the TCGA germline variant data and published summary statistics from genome-wide association studies (GWAS) (see Methods, Supplementary Table 1). The PRS validation results are summarized in Supplementary Table 2 (see Methods): all cancer PRS were positively associated with the corresponding cancer in the TCGA sample.

### Correlations with age at cancer diagnosis

We checked the correlations between somatic mutations by signature (and TSMC) and age at cancer diagnosis for each selected cancer type (Fig. 2). Consistent with previous study^5^, SBS1 and SBS5, the two clock-like signatures for which the numbers of mutations increase with age, showed strong positive correlations with age at diagnosis for most cancer types, though there exists heterogeneity across cancers. We also evaluated the correlation between the calculated germline PRS and age at diagnosis for each cancer type (Supplementary Fig. 1). Although none of these associations passed the Bonferroni-adjusted statistical significance threshold accounting for 322 tests (p = 1.55 × 10^−4^), the Spearman’s ρ with age for most cancer PRS were negative among the cases of that specific cancer, indicating that higher germline PRS of a cancer is associated with earlier diagnosis of that cancer.

**Fig. 2.**
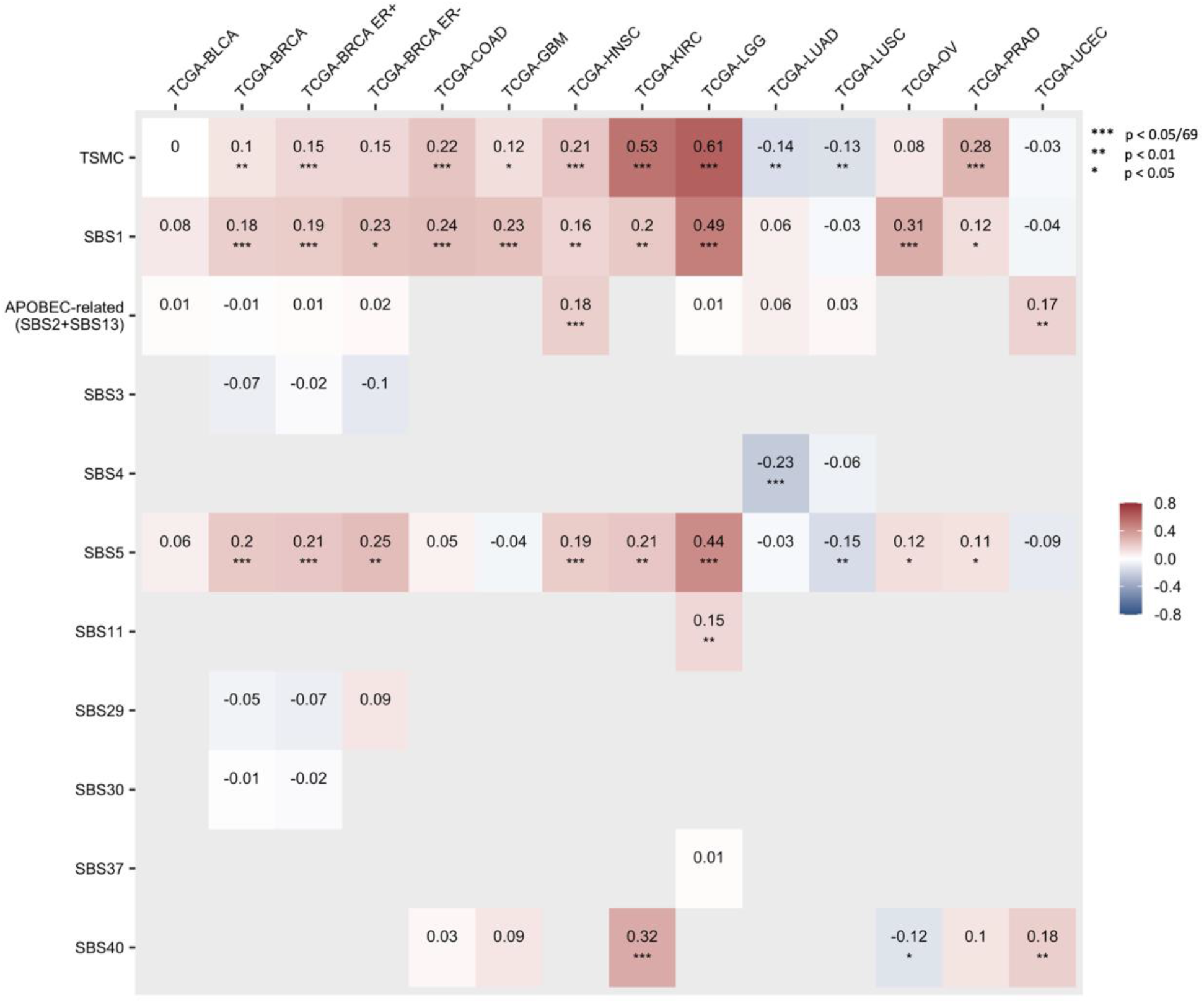
Correlations between somatic mutation counts and age at cancer diagnosis across cancers. Only the correlations with age at cancer diagnosis for those cancer-signature pairs included in the analyses are shown in this figure. Number in each cell and the cell color represent the Spearman correlation (ρ) between mutation counts of a SBS signature or TSMC (y-axis) and age at diagnosis in a cancer type (x-axis). Corrections passed the Bonferroni threshold (p < 0.05/69 = 0.00725) are marked with triple asterisk (***), correlations with p < 0.01 are marked with double asterisk (**), and correlations with p < 0.05 are marked with single asterisk (*).

### Associations between somatic mutations and germline PRS

We fit a zero-inflated negative binomial model, negative binomial model, or linear model regressing tumor somatic mutation counts on germline cancer or non-cancer PRS for each combination of cancer type (or subtype), SBS signature (or TSMC), and PRS, adjusting for sex (if applicable), age at cancer diagnosis (in years), and the top 10 genetic principal components (PCs). The p-value threshold for significance was 3.15 × 10^−5^, adjusting for multiple comparisons using Bonferroni correction (1,587 tests). We found 17 statistically significant associations (Table 1). Significant associations were found for prostate, breast, and endometrial cancers; most of them involved somatic mutation count of SBS1 (deamination of 5-methylcytosine) and the immune-mediated inflammatory diseases (IBD, CD, and UC) PRS. The association results across the 12 cancer types and two breast cancer subtypes for significant signature-PRS pairs are shown in Supplementary Fig. 2. Given that there was a strong positive correlation between number of mutations of SBS1 and age in breast cancer (Spearman’s ρ = 0.18, p = 2.50 × 10^−6^), a nominally significant correlation between SBS1 and age in prostate cancer (Spearman’s ρ = 0.12, p = 0.02), and a significant correlation between SBS40 and age in endometrial cancer (Spearman’s ρ = 0.18, p = 9.67 × 10^−4^), we further evaluated the associations of somatic mutations and PRS without adjusting for age at cancer diagnosis. There was no substantial change on the top findings, though we observed more significant associations with SBS1 and SBS5, which are likely to be driven by their correlations with age at diagnosis (Supplementary Table 3). The association between APOBEC-related signatures and breast cancer PRS in breast tumor is slightly above the Bonferroni threshold (p = 4.21 × 10^−5^) without adjusting for age.

**Table 1.**
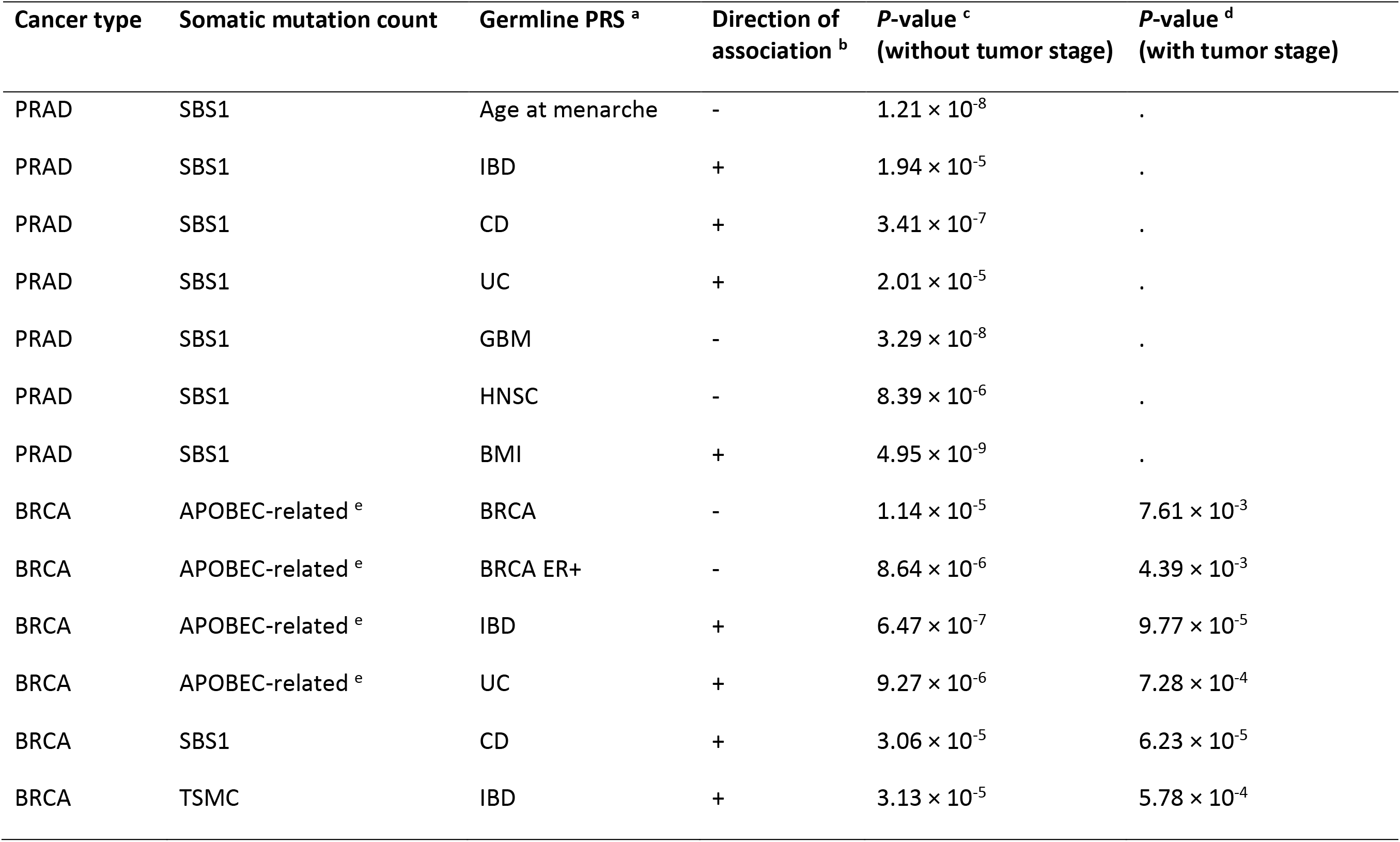
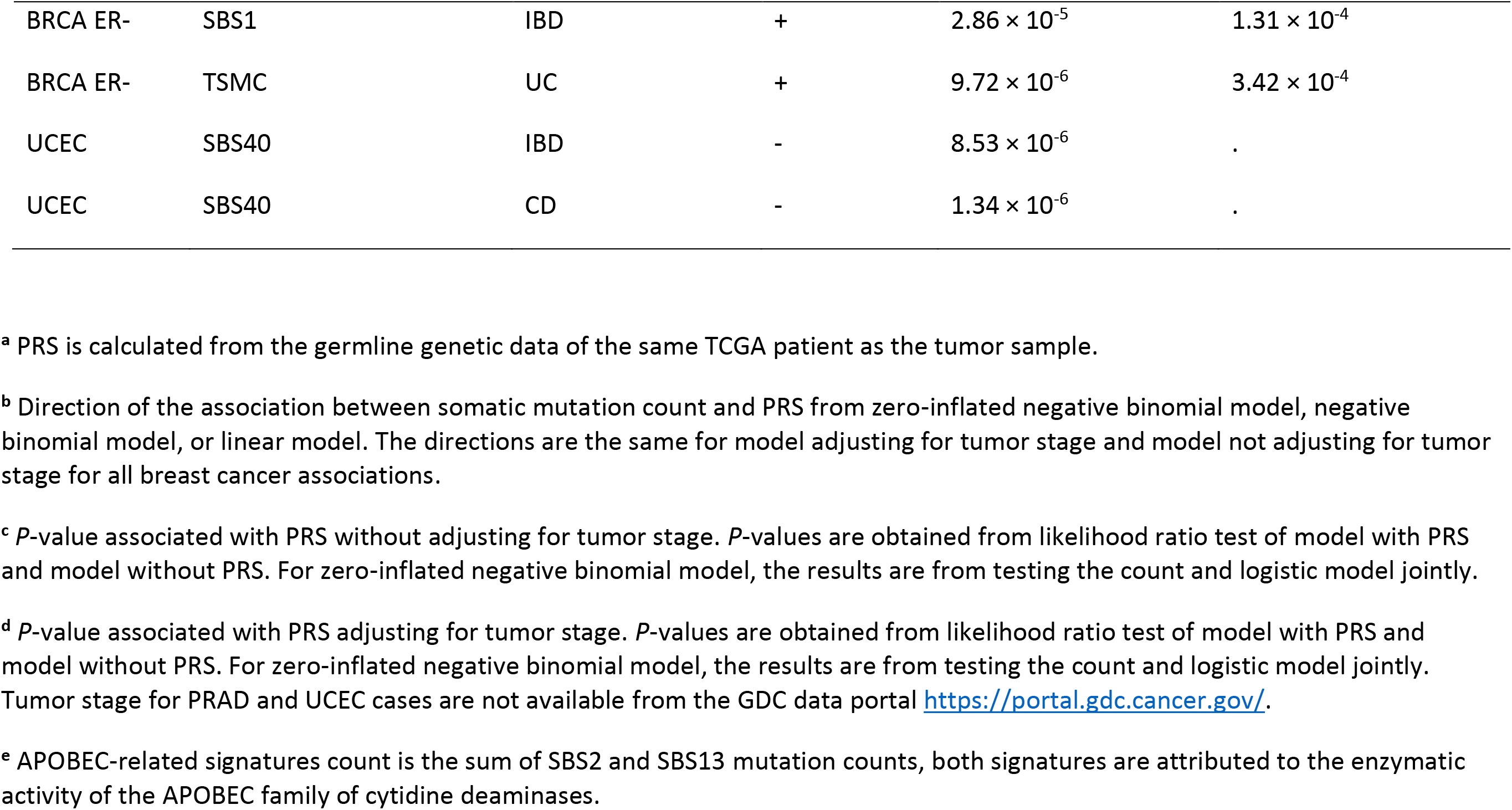
Significant associations between tumor somatic mutation counts and germline PRS.

| Cancer type | Somatic mutation count | Germline PRS <sup>a</sup> | Direction of association <sup>b</sup> | *P*-value <sup>c</sup><br>(without tumor stage) | *P*-value <sup>d</sup><br>(with tumor stage) |
| --- | --- | --- | --- | --- | --- |
| PRAD | SBS1 | Age at menarche | - | $1.21 \times 10^{-8}$ | . |
| PRAD | SBS1 | IBD | + | $1.94 \times 10^{-5}$ | . |
| PRAD | SBS1 | CD | + | $3.41 \times 10^{-7}$ | . |
| PRAD | SBS1 | UC | + | $2.01 \times 10^{-5}$ | . |
| PRAD | SBS1 | GBM | - | $3.29 \times 10^{-8}$ | . |
| PRAD | SBS1 | HNSC | - | $8.39 \times 10^{-6}$ | . |
| PRAD | SBS1 | BMI | + | $4.95 \times 10^{-9}$ | . |
| BRCA | APOBEC-related <sup>e</sup> | BRCA | - | $1.14 \times 10^{-5}$ | $7.61 \times 10^{-3}$ |
| BRCA | APOBEC-related <sup>e</sup> | BRCA ER+ | - | $8.64 \times 10^{-6}$ | $4.39 \times 10^{-3}$ |
| BRCA | APOBEC-related <sup>e</sup> | IBD | + | $6.47 \times 10^{-7}$ | $9.77 \times 10^{-5}$ |
| BRCA | APOBEC-related <sup>e</sup> | UC | + | $9.27 \times 10^{-6}$ | $7.28 \times 10^{-4}$ |
| BRCA | SBS1 | CD | + | $3.06 \times 10^{-5}$ | $6.23 \times 10^{-5}$ |
| BRCA | TSMC | IBD | + | $3.13 \times 10^{-5}$ | $5.78 \times 10^{-4}$ |
| BRCA ER- | SBS1 | IBD | + | $2.86 \times 10^{-5}$ | $1.31 \times 10^{-4}$ |
| BRCA ER- | TSMC | UC | + | $9.72 \times 10^{-6}$ | $3.42 \times 10^{-4}$ |
| UCEC | SBS40 | IBD | - | $8.53 \times 10^{-6}$ | . |
| UCEC | SBS40 | CD | - | $1.34 \times 10^{-6}$ | . |

<sup>a</sup> PRS is calculated from the germline genetic data of the same TCGA patient as the tumor sample.
<sup>b</sup> Direction of the association between somatic mutation count and PRS from zero-inflated negative binomial model, negative binomial model, or linear model. The directions are the same for model adjusting for tumor stage and model not adjusting for tumor stage for all breast cancer associations.
<sup>c</sup> *P*-value associated with PRS without adjusting for tumor stage. *P*-values are obtained from likelihood ratio test of model with PRS and model without PRS. For zero-inflated negative binomial model, the results are from testing the count and logistic model jointly.
<sup>d</sup> *P*-value associated with PRS adjusting for tumor stage. *P*-values are obtained from likelihood ratio test of model with PRS and model without PRS. For zero-inflated negative binomial model, the results are from testing the count and logistic model jointly. Tumor stage for PRAD and UCEC cases are not available from the GDC data portal https://portal.gdc.cancer.gov/.
<sup>e</sup> APOBEC-related signatures count is the sum of SBS2 and SBS13 mutation counts, both signatures are attributed to the enzymatic activity of the APOBEC family of cytidine deaminases.

There was no substantial change in the directions and effect sizes of the significant associations after adjusting for tumor stage where available (see Methods); the p-values for the breast cancer associations remained nominally significant after adjusting for stage (Table 1).

Consistent with a previous study^28^, which found a significant inverse association between breast cancer PRS and TSMC in breast tumor samples for ER+ patients, we found inverse associations between the somatic mutation counts of APOBEC-related signatures and breast cancer PRS (both overall and ER+ specific) in breast tumors. We further adjusted for the known germline APOBEC3 risk variant, rs17000526^29^, in the associations with the APOBEC-related signatures. This variant was not included in the calculation of any cancer or non-cancer PRS or in linkage disequilibrium (LD) (r^2^ > 0.1) with any PRS variants. There was no substantial change on the top associations with the APOBEC-related signatures. Consistent with a previous study^29^, we observed a significant association between rs17000526 and the APOBEC-related signatures in bladder tumors (regression coefficient = 0.21 for the rs17000526-A allele, p = 1.81 × 10^−4^).

This association was not significant in overall breast tumors. However, we did find a nominally significant association between rs17000526 and the APOBEC-related signatures in ER+ breast tumors (regression coefficient for the A allele = 0.10, p = 0.04).

Except for breast cancer, none of the somatic mutations and germline PRS associations for the same cancer type passed the significance threshold; the smallest p-value observed was for the positive association between prostate cancer PRS and SBS1 count in prostate tumor (p = 1.12 × 10^−3^). We did not see any significant association between mutation counts in lung tumor and cigarettes-per-day PRS (smallest p-values: p = 6.22 × 10^−3^ for TSMC in lung adenocarcinoma; p = 0.02 for SBS1 in lung squamous cell carcinoma). The smallest p-value for the cigarettes-per-day PRS was observed for SBS1 count in colorectal tumor (p = 1.70 × 10^−4^). Drinks-per-week and RA PRS were also not significantly associated with the mutation counts in any cancer type (smallest p-values: p = 2.26 × 10^−3^ for drinks-per-week PRS and SBS1 count in lung adenocarcinoma; p = 8.21 × 10^−4^ for RA and APOBEC-related signatures count in ER- breast tumor).

We further performed a meta-analysis on the association of SBS signatures (or TSMC) and germline PRS across cancers. The associations of SBS1 with CD PRS, SBS1 with bladder cancer PRS, and APOBEC-related signatures with IBD, CD, and UC PRS were significant using both fixed-effect and the Stouffer’s Z methods (p < 4.35 × 10^−4^ after Bonferroni correction for 115 tests) (Fig. 3). Overall, we found 17 associations that were statistically significant based on either the fixed-effect model or Stouffer’s method (Supplementary Table 4).

**Fig. 3.**
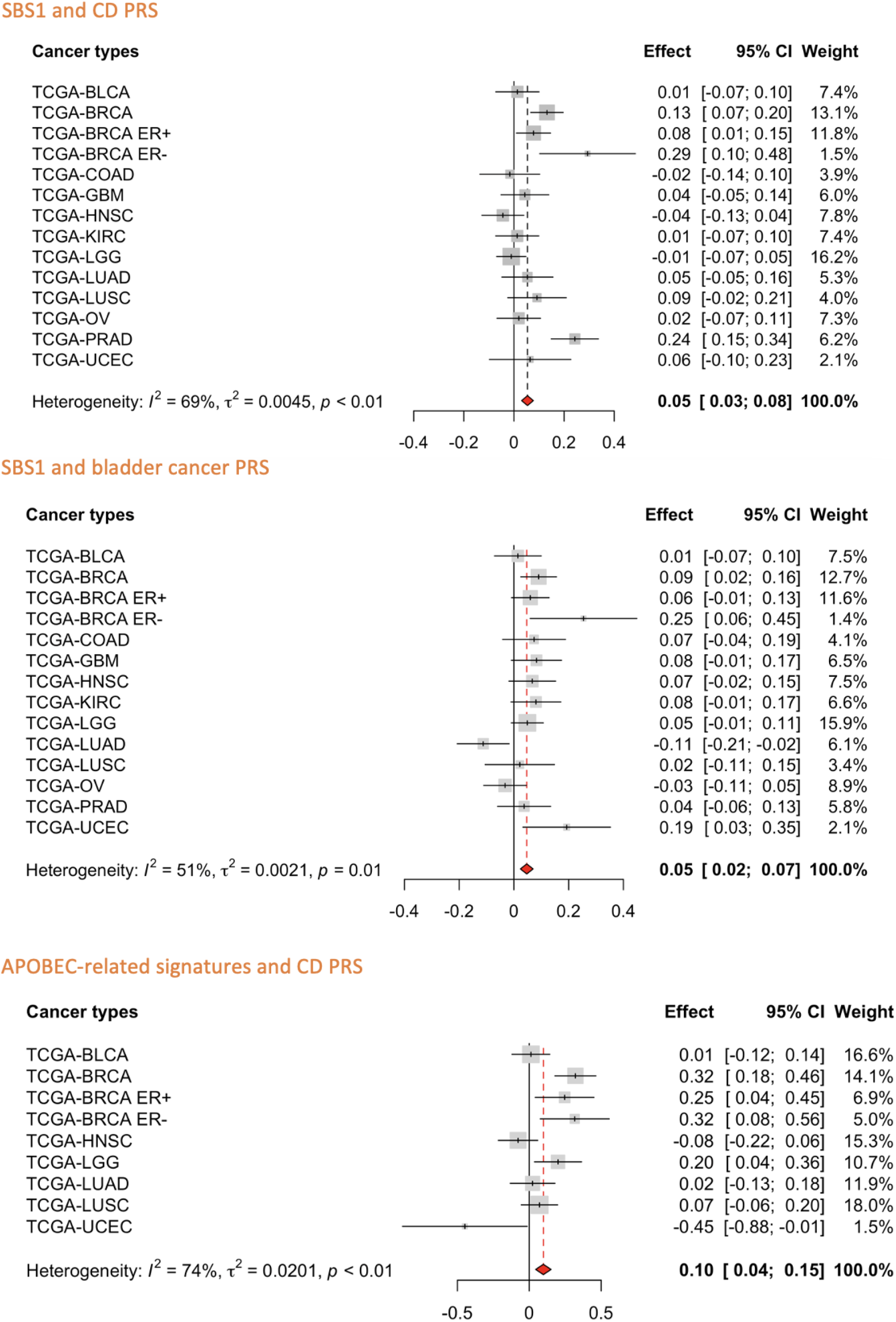
The associations between somatic mutation counts of SBS1 and CD PRS, SBS1 and bladder cancer PRS, and APOBEC-related signatures and CD PRS across cancers. The fixed-effect and Stouffer’s p-values for the association between SBS1 and CD PRS are p = 6.46 × 10^−6^ and p = 1.25 × 10^−5^; for the association between SBS1 and the bladder cancer PRS, they are p = 8.30 × 10^−5^and p = 4.01 × 10^−4^; and for the association between APOBEC-related signatures and CD PRS, they are 3.41 × 10^−4^ and p = 3.92 × 10^−4^). There are significant heterogeneities in the effect sizes for all three associations across cancers (p < 0.01). The effect sizes and 95% CI for PRS are plotted using grey squares and black horizontal lines. Size of the grey squares represents the weight in fixed-effect model for each cancer type. Red dashed line and red diamond represent the results from fixed-effect model.

**Fig. 4.**
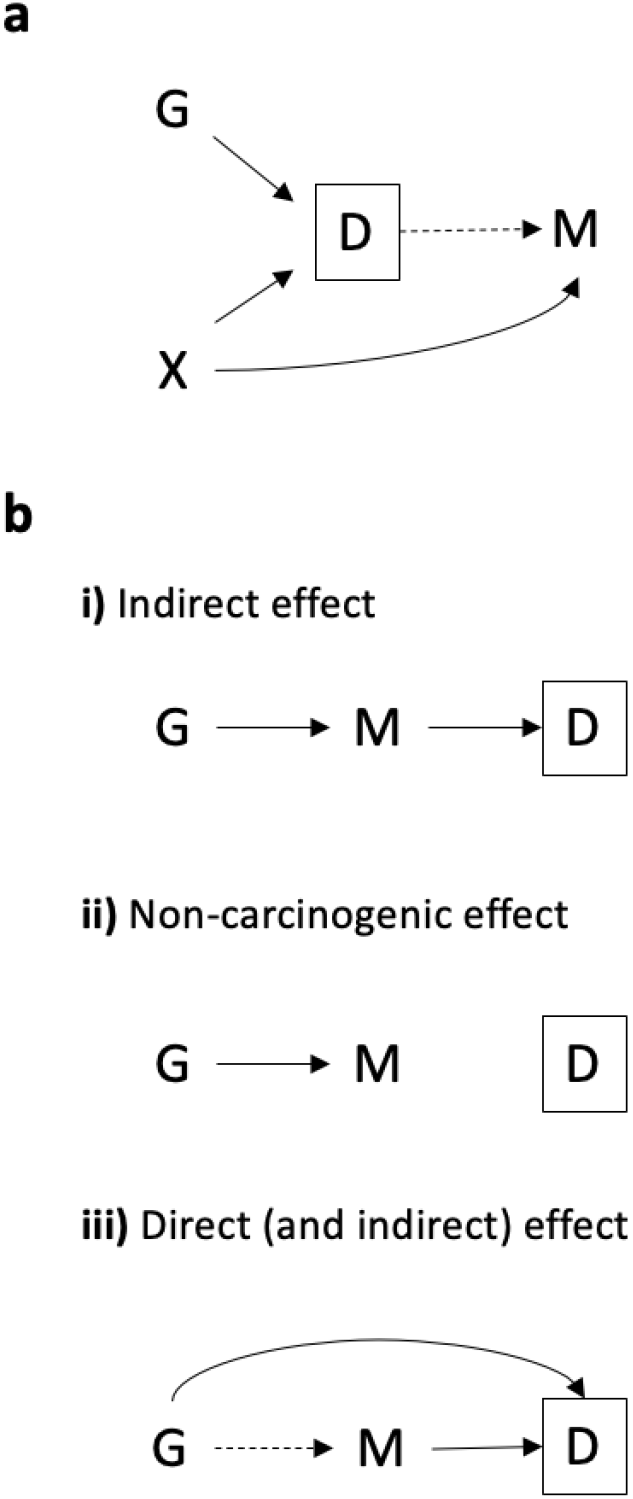
Hypothetical relationships between germline PRS, risk factor, mutational signature, and diagnosis of cancer. **a** Cancer PRS is associated with mutational signature due to collider bias. Cancer diagnosis (*D*) may or may not have an effect on somatic mutations of certain mutational signature (*M*). The cancer risk factor (*X*) is independent of cancer PRS (*G*) in the general population and is also associated with mutational signatures (*M*). Conditioning on diagnosis (*D*, i.e., studying cancer cases only) would induce collider bias on the relationship between cancer PRS (*G*) and mutational signature (*M*). **b** Three possible relationships between cancer or non-cancer PRS, mutational signature, and cancer diagnosis assuming no reverse causation. **(i) Indirect effect:** PRS (*G*) has an indirect effect on tumor development and diagnosis (*D*) through inducing somatic mutations of certain mutational signature (*M*); **(ii) Non-carcinogenic effect:** PRS (*G*) has an effect on inducing somatic mutations of certain mutational signature (*M*) but neither PRS (*G*) nor somatic mutations (*M*) has an effect on tumor development and diagnosis (*D*); **(c) Direct (and indirect) effect:** PRS (*G*) has a direct effect on tumor development and diagnosis (*D*) that not through the effect of somatic mutations (*M*) and may or may not have an indirect effect through somatic mutations (*M*). In this case, conditioning on diagnosis (*D*, i.e., studying cancer cases only) would induce collider bias on the relationship between PRS (*G*) and mutational signature (*M*).

## DISCUSSION

We performed a comprehensive analysis on the association between tumor somatic mutational profiles and germline PRS for various cancers and non-cancer traits, leveraging mutational signatures and germline variant data of 12 cancer types, as well as summary statistics from recent large GWAS. Our results demonstrate that there are robust associations between somatic mutational burden and germline PRS in human cancer. Some PRS can serve as proxies for exposures: linking mutational signatures to these exogenous and endogenous risk factors represented by PRS may suggest the etiology of associated mutational processes. Other PRS reflect the germline genetic contribution to cancer risk: studying their relationships with tumor somatic mutational burden may shed light on the underlying mechanism of cancer development that attributed to germline-somatic interactions.

For cancer PRS, the overall breast cancer PRS and ER+ specific PRS were both associated with the somatic mutation count of APOBEC-related signatures in breast tumor. These inverse relationships are consistent with a previous study by Zhu et al.^28^. Similarly, Qing et al.^32^ reported a significant inverse relationship between rare, high impact germline coding variants and somatic mutational burden in cancer hallmark genes among TCGA patients across age groups. It has been hypothesized that patients with higher germline variant burden tend to develop cancer at a younger age thus would have fewer acquired somatic mutations, whereas patients at a lower germline genetic risk may need longer time for cancer development, which is mainly driven by the accumulation of somatic mutations^28, 32^. However, we observed this inverse relationship for APOBEC-related signatures, which did not show clock-like behavior in previous studies^1, 5^. In our data, there was also no statistically significant correlation between mutation count of APOBEC-related signatures in breast cancer and age at diagnosis (Spearman’s ρ = −0.01, p = 0.73, Fig. 2). The p-value for this association increased after adjusting for tumor stage, but the direction of association did not change. This inverse relationship may indicate that breast tumors in women with low breast cancer PRS are predisposed to develop tumors with higher APOBEC mutation counts (due to yet-to-be-determined biological mechanisms), but it may also be a result of collider bias (Fig. 3a): if other breast cancer risk factors that are independent of the breast cancer PRS in the general population are associated with higher APOBEC mutation counts, then the breast cancer PRS and APOBEC mutation counts could be negatively correlated among cases. Few studies have directly assessed the associations between established cancer risk factors and SBS signatures. A recent study reported no impact of BMI, cigarette smoking, and alcohol consumption on APOBEC signatures in TCGA^33^.

In another simple setting assuming no reverse causation, there might be three possible causal relationships between cancer or non-cancer PRS, mutational signature, and cancer diagnosis given that a significant association between a germline PRS and a mutational signature is observed (Fig. 3b). Collider bias would also arise if both the PRS and mutational signature have a direct effect (or effect through other pathways) on tumor development and cancer diagnosis. If that is the case, a non-causal association between germline PRS and mutational signature would be observed if we only include cancer cases in the study. This scenario is plausible for breast cancer, as many breast cancer susceptibility variants identified from GWAS have been shown to directly code or regulate cancer susceptibility genes (e.g., missense variant rs35383942 is in *PHLDA3*, which encodes a p53-regulated repressor of Akt^34^), thus have effects on breast cancer development through other functional pathways that are not mediated by the acquirement of somatic mutations^35, 36, 37, 38^. To avoid the potential collider biases, somatic mutation profiling needs to be performed on both tumor and normal tissue at the cancer sites of interest and the sampling of study population should be independent of cancer status. The Human Tumor Atlas Network^39^, which seeks to construct comprehensive atlases of molecular and cellular features of cancers, starting from precancerous lesions to advanced disease, could address these questions. In addition to the breast cancer associations, inverse relationships were also observed for other two cancer PRS associations (head and neck cancer PRS and glioblastoma PRS with SBS1 in prostate tumor), either adjusting or not adjusting for age at diagnosis, which may also be explained by the collider bias described above.

For non-cancer PRS, germline PRS of age at menarche was found to be significantly associated with SBS1 count in prostate tumor. Specifically, we found that prostate cancer patients with higher age at menarche PRS, a surrogate for later menarche, tend to have fewer SBS1 mutations. Although males do not exhibit menarche, a previous study reported a strong genetic correlation (r_g_ = 0.74) between female and male puberty timing, represented by age at menarche and age at voice breaking, respectively^40^. Therefore, we hypothesize that the effect of hormonal factor PRS on the number of somatic mutations in prostate tumors may be explained by the shared regulatory mechanism of puberty timing in men and women. Prostate cancer has long been recognized as a hormone-related cancer, traditional epidemiological studies have reported associations between various hormone-related markers (e.g., Insulin-like growth factor 1 (IGF-1), testosterone, sex hormone-binding globulin, etc.) and the risk of prostate cancer^41, 42, 43, 44^. Mendelian randomization studies also reported a consistent protective effect of later pubertal development on prostate cancer risk^40, 45^. All these suggest an etiological relevance between the timing of puberty and incidence of prostate cancer involving shared effect of hormonal factors. Previous findings highlighted the roles of androgen and IGF-1, of which the circulating levels increase dramatically during puberty, in prostate carcinogenesis^45, 46, 47, 48, 49^. They proposed that the concentrations of these hormones during this susceptibility window when the luminal cells start to appear and the prostate becomes mature may have an impact on prostate cancer risk in later life^50, 51, 52, 53^. Our finding suggests a potential pathway through the hormone-related markers of puberty timing on the accumulation of SBS1 mutations in prostate tissue, which may drive tumor development.

Inflammatory bowel disease PRS were found to be associated with somatic mutational profiles in multiple hormone-related cancers (breast, endometrial, and prostate cancer). Previous studies have established CD and UC as risk factors for overall cancer^54, 55, 56, 57, 58^, but whether these associations are driven by shared genetic susceptibility or other common lifestyle and environmental factors remains unanswered by these epidemiological studies. In the present work, we found robust positive associations between IBD PRS (IBD, UC, and CD) and somatic mutations in breast cancer. Previous evidence has shown an increased risk of breast cancer among UC and CD patients, and first-degree relatives of CD patients^57, 59, 60^. Several potential mechanisms have been proposed. Interleukin-1 (IL-1) polymorphism has been linked the risk of many diseases, including breast cancer and IBD^61, 62, 63, 64, 65, 66^. A recent study^67^ identified 53 common differentially expressed genes for breast cancer and CD compared with controls, three major hub genes, *CXCL8*, *IL-1β* and *PTGS2*, and the shared IL-17 and NF-κB signaling pathways. These suggest that the elevation of cytokines may play a common role in the pathogenesis of CD and breast cancer. Hovde et al.^57^ also proposed that the downregulation of breast cancer resistance protein in UC patients may be related to breast cancer etiology. In addition to breast cancer, we also observed positive relationships between IBD PRS (IBD, UC, and CD) and somatic mutation count of SBS1 in prostate tumor. Moreover, the association between CD and SBS1 is broadly significant across the 12 cancer types in our study. Recent meta-analyses and a large cohort study all conclude that patients with IBD have an increased risk of prostate cancer^68, 69, 70^, though the previous findings were inconclusive^71, 72, 73, 74, 75, 76^. Previous studies have proposed that shared risk alleles may partially explain this association^68, 75, 77^. Folate hydrolase 1 (FOLH1) or prostate-specific membrane antigen (PSMA) is highly expressed in both IBD and prostate cancer^78, 79, 80^. Studies have reported that Inhibition of FOLH1/PSMA ameliorates IBD symptom in mice models^78^ and also leads to tumor regression in preclinical models^81^. Our results suggest there is a link between inherited genetic variants that contribute to the development of inflammatory bowel disease and acquired somatic mutations in these hormonal-related cancers. Future studies need to further investigate the mechanism underlying the associations between specific PRS and specific SBS signatures, especially between CD and SBS1. One direction might be looking at the associations between these mutational signatures and the measured immune-related markers in TCGA. This may also provide novel insight into the mechanism underlying the association between TMB and benefit from immunotherapy.

Interestingly, we found a significant inverse association between IBD PRS (IBD and CD) and SBS40 count in endometrial cancer. There are limited studies looking at the association between IBD and risk of endometrial cancer. A meta-analysis reported an inverse relationship, but it is not statistically significant^71^. Our results may suggest potential mechanisms underlying SBS40 through immune-related processes.

A significant positive association was observed between BMI PRS and SBS1 count in prostate cancer. Obesity has been established as an independent risk factor associated with the risk of advanced or fatal prostate cancer^82, 83, 84^. Studies also have shown that high BMI is associated with a decreased risk of low-grade prostate cancer^82, 83, 85, 86, 87^. However, these associations may be due to diagnostic bias. Men with higher BMI tend to have lower prostate-specific antigen levels^88, 89^, which make them less likely to have a biopsy^90^, and larger prostate volumes, which makes it harder to find the tumor from biopsy^91^. Mendelian randomization studies have found little evidence for a causal relationship between BMI and prostate cancer incidence^92, 93^. Therefore, it is likely that higher BMI PRS is associated with older age at diagnosis, and since SBS1 counts increase with age, people with higher BMI PRS would have higher SBS1 counts as well. We did not adjust for tumor stage or grade for prostate cancer cases, as it was unavailable. We adjusted for age at diagnosis, and the BMI PRS is not significantly association age in prostate cancers in our study (Supplementary Fig. 1), but we still observed this strong positive association. Future studies need to further investigate this association with SBS1 incorporating tumor stage and grade information, and carefully control for the potential biases.

Our study has limitations. The calculated PRS only explains a small proportion of the heritability thus may not fully represent the genetic variant burden underlying the trait. Although we adjusted for tumor stage for most cancer types, we did not adjust for or perform subgroup analysis by tumor grade, as this information is not available for the selected TCGA patients. Bonferroni correction for multiple testing is conservative. We expect to identify more significant associations with a larger sample size. There are several strengths of our study. We performed a comprehensive pan-cancer analysis on the relationship between 14 cancer PRS and 9 non-cancer PRS, and TSMC as well as the number of mutations attributed to 10 SBS signatures across 12 cancer types. We used PRS as genetic instruments for endogenous and exogenous exposures to avoid bias from confounding and reverse causation in traditional observational studies. Also, we have sufficient sample size for detecting associations of similar strength as previous study^28^ in each cancer type with a high power, which may not be able to achieve using exposure data that usually have missing metadata. We controlled for the impact of age at diagnosis and tumor stage on the germline-somatic associations. Although PRS serves as a good proxy for exposure, it would still be useful to collect epidemiological data on exposures in future studies to confirm our findings and to better understand the role of germline variant burden underlying these associations. Future studies can also look at the association between signature-specific mutation count and immune features, hormone-related markers, and expression levels of cancer susceptibility genes to further investigate the underlying mechanisms.

In conclusion, our findings indicate that there are robust associations between somatic mutational profiles and germline PRS in human cancer. Our results demonstrate evidence for germline-somatic associations between inflammatory bowel disease PRS and somatic mutations (SBS1, APOCEC-related signatures, SBS40, and TSMC) in breast cancer, endometrial cancer, and prostate cancer, and between age at menarche PRS and somatic mutations (SBS1) in prostate cancer. Our results are relevant to the etiology of mutational signatures and the underlying biological mechanisms of cancer development.

## METHODS

### Study population

We selected cancer types based on total number of cases in TCGA and the availability of large GWAS for calculating PRS. Twelve cancer types with WES data available on more than 300 primary tumor samples were selected to ensure 80% power (at a type I error rate of 5%) to detect an association between PRS and TSMC of similar or greater magnitude as that previously reported between rs2588809 and TMB^28^ (Supplementary Fig. 3). Data were obtained for all TCGA patients of European ancestry from these 12 cancer types. We further restricted sample type to primary tumor and excluded male cases for breast cancer. Age at diagnosis, sex, and tumor stage were retrieved from the GDC data portal https://portal.gdc.cancer.gov/. Tumor stage was not available for prostate cancer, endometrial cancer, lower grade glioma, glioblastoma, and ovarian cancer. Samples with missing age at diagnosis, tumor stage (if available), and sex information were further excluded.

### Mutational signatures

Mutational signatures were identified from TCGA WES data using methods based on nonnegative matrix factorization^1^. We obtained the mutation counts for 54 distinct SBS signatures for each selected tumor sample from the ICGC data portal https://dcc.icgc.org/releases/PCAWG. In addition, we created a new signature variable called APOBEC-related signatures by summing up the somatic mutation counts of SBS2 and SBS13 (both attributed to APOBEC activity). For each cancer type, we analyzed those SBS signatures if there was at least one cancer type where both more than 20 samples and more than 20% of the samples in the cancer type had mutations attributed to that signature. In addition to the SBS signature-specific mutations, we also retrieved TSMC for each sample using maftools R package^94^.

### Germline variant data

Raw germline single nucleotide polymorphism (SNP) array data were downloaded from the GDC Legacy Archive https://portal.gdc.cancer.gov/legacy-archive. Genotype quality control (QC) was applied to remove variants with >5% missing calls, or Hardy-Weinberg equilibrium p < 5×10^−6^, or minor allele frequency (MAF) <1%. To restrict to individuals of European ancestry, genetic PCs were computed using post-QC, LD pruned variants and any outlier sample >6 standard deviations away from the mean along either of the top 2 PCs was removed. All remaining samples and genotypes were then imputed to the Haplotype Reference Consortium reference panel^95^.

### Calculation of PRS

The PRS of a trait for subject *i* was calculated as,

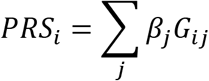

where the weight *β_j_* was the log odds ratio (or the beta coefficient for continuous traits) of the trait comparing effect allele to other allele at SNP *j*, and *G_ij_* was the expected number of effect allele at SNP *j* for subject *i* (allele dosage). The lists of SNPs for constructing PRS were obtained from published GWAS summary statistics (Supplementary Table 1).

For cancer PRS, the list of SNPs and corresponding weights were obtained from one of the three sources: *(i)* GWAS or PRS paper; *(ii)* NHGRI-EBI GWAS Catalog^96^; and *(iii)* Cancer PRSweb^97^. For non-cancer PRS, the summary statistics were all from GWAS or PRS papers. Sources of GWAS summary statistics for each trait are summarized in Supplementary Table 1.

We filtered the SNP list for each trait using different strategies. For those from GWAS Catalog, we removed results from cross-cancer, subgroup, or interaction analysis, and restricted to European ancestry studies with a minimum of 2,000 cases and 2,000 controls. We removed SNPs with p-value above the genome-wide significance threshold (p > 5×10^−8^) for those from GWAS Catalog or GWAS papers. For SNPs from Cancer PRSweb, we used the p-values threshold with the best performance as evaluated by Nagelkerke’s pseudo-R^2^. We did not additionally filter by p-values nor perform LD clumping on SNPs from PRS papers. LD clumping was performed on SNP lists from GWAS paper and GWAS Catalog: we removed SNPs with MAF < 1% or in LD (r^2^ > 0.1) with SNPs of smaller p-value. We used PLINK 2.0^98^ to calculate PRS for each trait and subject from the final list of SNPs.

### Validation of PRS

We evaluated the ability of the calculated PRS to discriminate between specific cancer cases and controls. For each cancer type and PRS, an unadjusted logistic model was fit by treating all patients of that cancer type as cases and a randomly selected subset (with the same size as cases) of other cancer cases as controls. The performance of PRS was assessed by area under the receiver operating characteristic curve (Supplementary Table 2). In addition to the 12 cancer types and two subtypes, we also evaluated the performance of PRS on discriminating lung cancer cases combined (lung adenocarcinoma and squamous cell carcinoma) and glioma cases combined (lower grade glioma and glioblastoma). For those cancer types with subtypes or other closely related types (e.g., breast cancer), none of the patients of the other subtypes (or related types) was included as controls when treating one subtype (or type) as cases.

### Association between somatic mutation count and PRS

TSMC were Winsorized to 98%, where the counts below the first percentile were set to the first percentile and the counts above the 99th percentile were set to the 99th percentile. PRS was standardized to have mean zero and unit standard deviation before running each model. For each cancer type and SBS signature (or TSMC), we first fit a zero-inflated negative binomial model if the proportion of zero-count samples is greater than 10%, otherwise we fit a negative binomial model directly. If the zero-inflated negative binomial model failed to converge and the proportion of zero-count samples is less than 50% (if greater than or equal to 50%, all models were considered as failed), we tried negative binomial model. If both two models failed, we transformed the mutation count to log_10_ (count + 1) and fit the linear model. In each attempt, we ran both model with and without PRS (models were considered as failed if either model failed) and obtained p-value from likelihood ratio test on these two models. We performed a fixed-effect meta-analysis of the associations between TSMC or each SBS signature and each PRS across cancers. Stouffer’s Z-score method was also used to combining the individual p-values from each cancer type. The Z-score for the overall meta-analysis is:

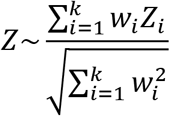

where *Z_i_* is the Z-score for cancer type *i*, *w_i_* is the sample size of cancer type *i*, *k* is the total number of cancer types and subtypes. We used Bonferroni correction to adjust for multiple comparisons of 115 tests, which is the total number of signature-PRS pairs, excluding those only available for breast cancer (overall, ER+, and ER-) and lung cancer (adenocarcinoma and squamous cell carcinoma).

### Sensitivity analysis

To assess the impact of age at cancer diagnosis on the association of germline PRS and somatic mutational burden, we ran the association tests without adjusting for age at diagnosis (same model types as adjusting for age). Spearman correlations were calculated for *(i)* somatic mutation count and age at diagnosis for each cancer type and signature. We used Bonferroni correction accounting for 69 tests, which is the total number of cancer-signature pairs included in the analyses; and *(ii)* PRS and age at diagnosis for each cancer type. We used Bonferroni correction accounting for 322 tests, which is the total number of cancer-PRS pairs. Tumor stage is only available for some cancers; we re-ran the association tests adjusting for tumor stage for those cancer types (same model types as not adjusting for tumor stage). We further adjusted for the known germline APOBEC3 risk variant, rs17000526, in the associations with the APOBEC-related signatures (same model types as not adjusting for rs17000526).

## DATA AVAILABILITY

The TCGA germline variants data are available at https://portal.gdc.cancer.gov/legacy-archive. The TCGA somatic mutations and mutational signatures data are available from the maftools R package^94^ and at https://dcc.icgc.org/releases/PCAWG. The TCGA clinical data are available at https://portal.gdc.cancer.gov/. Other relevant data are available from the authors upon request.

## ACKNOWLEDGMENT

This work was supported by grants from the U.S. National Institutes of Health (U01CA194393 and R01CA227237). The results published here are in whole or part based upon data generated by the TCGA Research Network: https://www.cancer.gov/tcga.

## AUTHOR CONTRIBUTIONS

P.K., A.G., and Y.J.H. designed the study. P.K. and Y.L. developed the analysis plan. L.B.A. prepared the somatic data of TCGA. A.G. prepared the germline data of TCGA. Y.L. performed the analyses. P.K. and Y.L wrote the manuscript. A.G., Y.J.H., and L.B.A. reviewed the manuscript.

## COMPETING INTERESTS

The authors declare no competing interests.

## MATERIALS & CORRESPONDENCE

Correspondence and requests for materials should be addressed to P.K.

## Notes

### Competing Interest Statement

The authors have declared no competing interest.

