## Supplementary Information for "Somatic mutational profiles and germline polygenic risk scores in human cancer"

### SUPPLEMENTARY TABLES

**Supplementary Table 1.** Sources of GWAS summary statistics for calculating PRS

**Supplementary Table 2.** PRS validation results

**Supplementary Table 3.** Significant results without adjusting for age at diagnosis ($p < 3.15\times10^{-5}$)

**Supplementary Table 4.** Significant results from meta-analyses ($p < 4.35 \times 10^{-4}$)

### SUPPLEMENTARY FIGURES

**Supplementary Fig. 1.** Correlations between germline PRS and age at cancer diagnosis across cancers

**Supplementary Fig. 2.** Significant associations between SBS signatures (or TSMC) and germline PRS across cancers.

**Supplementary Fig. 3.** Power as a function of proportion of variance in TSMC explained by PRS for various sample sizes

**Supplementary Table 1.** Sources of GWAS summary statistics for calculating PRS

| | Phenotype | Number of SNPs | GWAS Source | PMID/EFO ID/phecode[a] | Access/Publication Date |
|---|---|---|---|---|---|
| **Cancer PRS** | BLCA | 15 | Cancer PRSweb[1, 2, 3, 4, 5, 6, 7, 8] | 189.2 (cancer of bladder) | 4/23/2020 |
| | BRCA | 313 | PRS paper[9] | 30554720 | Jan-2019 |
| | BRCA ER+ | 313 | PRS paper[9] | 30554720 | Jan-2019 |
| | BRCA ER- | 313 | PRS paper[9] | 30554720 | Jan-2019 |
| | COAD | 87 | Cancer PRSweb[1, 10] | 153 (colorectal cancer) | 4/23/2020 |
| | GBM | 5 | Cancer PRSweb[1, 11, 12] | 191.11 (cancer of brain) | 4/23/2020 |
| | HNSC | 8 | GWAS paper[13] | 27749845 | Dec-2016 |
| | KIRC | 8 | GWAS Catalog[14, 15, 16] | EFO_0000681 (renal cell carcinoma) | 5/27/2020 |
| | LGG | 19 | GWAS Catalog[12, 14] | EFO_0005543 (glioma) | 5/27/2020 |
| | LUAD | 15 | GWAS Catalog[14, 17, 18] | EFO_0000571 (lung adenocarcinoma) | 5/27/2020 |
| | LUSC | 6 | GWAS Catalog[14, 18] | EFO_0000708 (squamous cell lung carcinoma) | 5/27/2020 |
| | OV | 21 | Cancer PRSweb[1, 19] | 184.11 (malignant neoplasm of ovary) | 4/23/2020 |

| | | | | | |
|---|---|---|---|---|---|
| | PRAD | 980 | Cancer PRSweb[1, 20] | 185 (cancer of prostate) | 4/23/2020 |
| | UCEC | 18 | Cancer PRSweb[1, 21, 22, 23] | 182 (malignant neoplasm of uterus) | 4/23/2020 |
| **Non-cancer PRS** | Age at menarche | 376 | GWAS paper[24] | 28436984 | Jun-2017 |
| | Age at natural menopause | 46 | PRS paper[25] | 27760082 | Feb-2017 |
| | BMI | 74 | GWAS paper[26] | 27427428 | Jun-2016 |
| | Cigarettes per day | 46 | GWAS paper[27] | 30643251 | Feb-2019 |
| | Drink per week | 90 | GWAS paper[27] | 30643251 | Feb-2019 |
| | IBD | 155 | GWAS paper[28] | 26192919 | Sep-2019 |
| | UC | 86 | GWAS paper[28] | 26192919 | Sep-2019 |
| | CD | 140 | GWAS paper[28] | 26192919 | Sep-2019 |
| | RA | 61 | GWAS paper[29] | 24390342 | Feb-2014 |

[a] PMID is listed if from PRS/GWAS paper; EFO ID is listed if from GWAS Catalog; phecode is listed if from Cancer PRSweb.

**Supplementary Table 2.** PRS validation results

| **Parallel-trait (cancer PRS validation) analyses** | | | | |
|---|---|---|---|---|
| **Cancer type** | **PRS** | **log OR** | ***P*-value** | **AUC** |
| BLCA | BLCA | 0.73 | 2.30E-03 | 0.56 |
| BRCA | BRCA | 0.75 | 4.64E-13 | 0.61 |
| BRCA ER+ | BRCA ER+ | 0.78 | 2.18E-12 | 0.62 |
| BRCA ER- | BRCA ER- | 0.66 | 6.09E-03 | 0.6 |
| COAD | COAD | 1.1 | 1.15E-08 | 0.63 |
| GBM | GBM | 1.07 | 2.67E-10 | 0.64 |
| HNSC | HNSC | 0.85 | 1.07E-04 | 0.58 |
| KIRC | KIRC | 0.85 | 2.82E-03 | 0.56 |
| LGG | LGG | 0.95 | 3.79E-14 | 0.65 |
| LUAD | LUAD | 1.15 | 6.02E-08 | 0.61 |
| LUSC | LUSC | 0.64 | 8.44E-03 | 0.54 |
| OV | OV | 1.12 | 9.03E-11 | 0.62 |
| PRAD | PRAD | 0.28 | 1.32E-04 | 0.58 |
| UCEC | UCEC | 1.1 | 7.48E-06 | 0.6 |
| **Cross-trait analyses** ($p < 1.36 \times 10^{-4}$)[a] | | | | |
| **Cancer type** | **PRS** | **log OR** | ***P*-value** | **AUC** |
| BRCA | BRCA ER+ | 0.71 | 3.48E-13 | 0.61 |
| BRCA | BRCA ER- | 0.57 | 4.61E-08 | 0.57 |
| BRCA ER+ | BRCA | 0.81 | 6.38E-12 | 0.62 |
| BRCA ER+ | BRCA ER- | 0.56 | 2.62E-06 | 0.57 |
| GBM | LGG | 0.55 | 6.40E-05 | 0.59 |
| GBM+LGG | GBM | 0.67 | 3.76E-10 | 0.59 |
| GBM+LGG | LGG | 0.71 | 2.01E-15 | 0.62 |
| HNSC | Drink per week | 4.16 | 3.66E-06 | 0.59 |
| LGG | GBM | 0.59 | 7.41E-05 | 0.57 |
| LUAD+LUSC | LUAD | 0.62 | 1.55E-05 | 0.56 |
| LUAD+LUSC | LUSC | 0.81 | 3.58E-06 | 0.56 |

[a] Bonferroni-adjusted statistical significance threshold accounting for 368 tests.

**Supplementary Table 3.** Significant results without adjusting for age at diagnosis ($p < 3.15\times10^{-5}$)

| **Cancer type** | **Somatic mutation count** | **Germline PRS** | **Direction of association** | ***P*-value** |
|---|---|---|---|---|
| PRAD | SBS1 | Age at menarche | - | 1.02E-08 |
| PRAD | SBS1 | IBD | + | 7.14E-06 |
| PRAD | SBS1 | CD | + | 1.34E-07 |
| PRAD | SBS1 | UC | + | 1.04E-05 |
| PRAD | SBS1 | GBM | - | 4.32E-08 |
| PRAD | SBS1 | HNSC | - | 2.58E-05 |
| PRAD | SBS1 | BMI | + | 1.25E-08 |
| PRAD | SBS5 | Age at natural menopause | - | 1.39E-06 |
| PRAD | SBS5 | BRCA ER+ | + | 5.32E-06 |
| BRCA | APOBEC-related | IBD | + | 4.45E-06 |
| BRCA | SBS1 | IBD | + | 2.2E-06 |
| BRCA | SBS1 | CD | + | 1.92E-06 |
| BRCA | TSMC | IBD | + | 1.38E-05 |
| BRCA | SBS1 | LUAD | + | 9.08E-06 |
| BRCA ER- | SBS1 | IBD | + | 2.76E-05 |
| BRCA | SBS5 | IBD | + | 2.75E-05 |
| BRCA ER- | TSMC | UC | + | 2.03E-05 |
| UCEC | SBS40 | IBD | - | 6.15E-07 |
| UCEC | SBS40 | CD | - | 6.08E-08 |
| GBM | SBS1 | OV | + | 6.56E-06 |

**Supplementary Table 4.** Significant results from meta-analyses ($p < 4.35 \times 10^{-4}$)

| Somatic mutation count | Germline PRS | Beta | SE | *P*-value (fixed-effect) | *P*-value (Cochran's Q) | $I^2$ | *P*-value (Stouffer's) |
|---|---|---|---|---|---|---|---|
| SBS1 | Age at menarche | 0.03 | 0.01 | 1.23E-02 | 1.57E-06 | 0.75 | 4.55E-05 |
| SBS1 | CD | 0.05 | 0.01 | 6.46E-06 | 7.65E-05 | 0.69 | 1.25E-05 |
| SBS1 | HNSC | -0.03 | 0.01 | 3.21E-02 | 3.50E-08 | 0.79 | 4.38E-06 |
| SBS1 | IBD | 0.05 | 0.01 | 1.87E-05 | 1.35E-04 | 0.68 | 4.95E-04 |
| SBS1 | LUAD | 0.03 | 0.01 | 9.00E-03 | 2.15E-04 | 0.66 | 4.94E-05 |
| SBS1 | LGG | -0.03 | 0.01 | 7.50E-03 | 4.59E-03 | 0.57 | 2.28E-04 |
| SBS1 | LUSC | 0.02 | 0.01 | 1.33E-01 | 3.49E-04 | 0.65 | 8.51E-05 |
| SBS1 | BLCA | 0.05 | 0.01 | 8.30E-05 | 1.50E-02 | 0.51 | 4.01E-04 |
| APOBEC-related | Age at menarche | -0.04 | 0.03 | 1.77E-01 | 1.96E-07 | 0.83 | 4.34E-07 |
| APOBEC-related | IBD | 0.13 | 0.03 | 2.31E-06 | 1.88E-04 | 0.74 | 5.49E-05 |
| APOBEC-related | CD | 0.10 | 0.03 | 3.41E-04 | 1.31E-04 | 0.74 | 3.92E-04 |
| APOBEC-related | UC | 0.10 | 0.03 | 2.40E-04 | 2.77E-05 | 0.77 | 1.57E-05 |
| APOBEC-related | BRCA | -0.07 | 0.03 | 1.40E-02 | 4.92E-08 | 0.84 | 9.72E-06 |
| APOBEC-related | BRCA ER+ | -0.06 | 0.03 | 2.48E-02 | 2.43E-08 | 0.84 | 7.85E-06 |
| APOBEC-related | BRCA ER- | -0.09 | 0.03 | 1.55E-03 | 2.34E-04 | 0.73 | 3.67E-05 |
| APOBEC-related | UCEC | -0.06 | 0.03 | 1.67E-02 | 3.19E-09 | 0.86 | 1.64E-08 |
| SBS5 | GBM | -0.02 | 0.01 | 6.37E-02 | 2.50E-04 | 0.66 | 1.53E-04 |

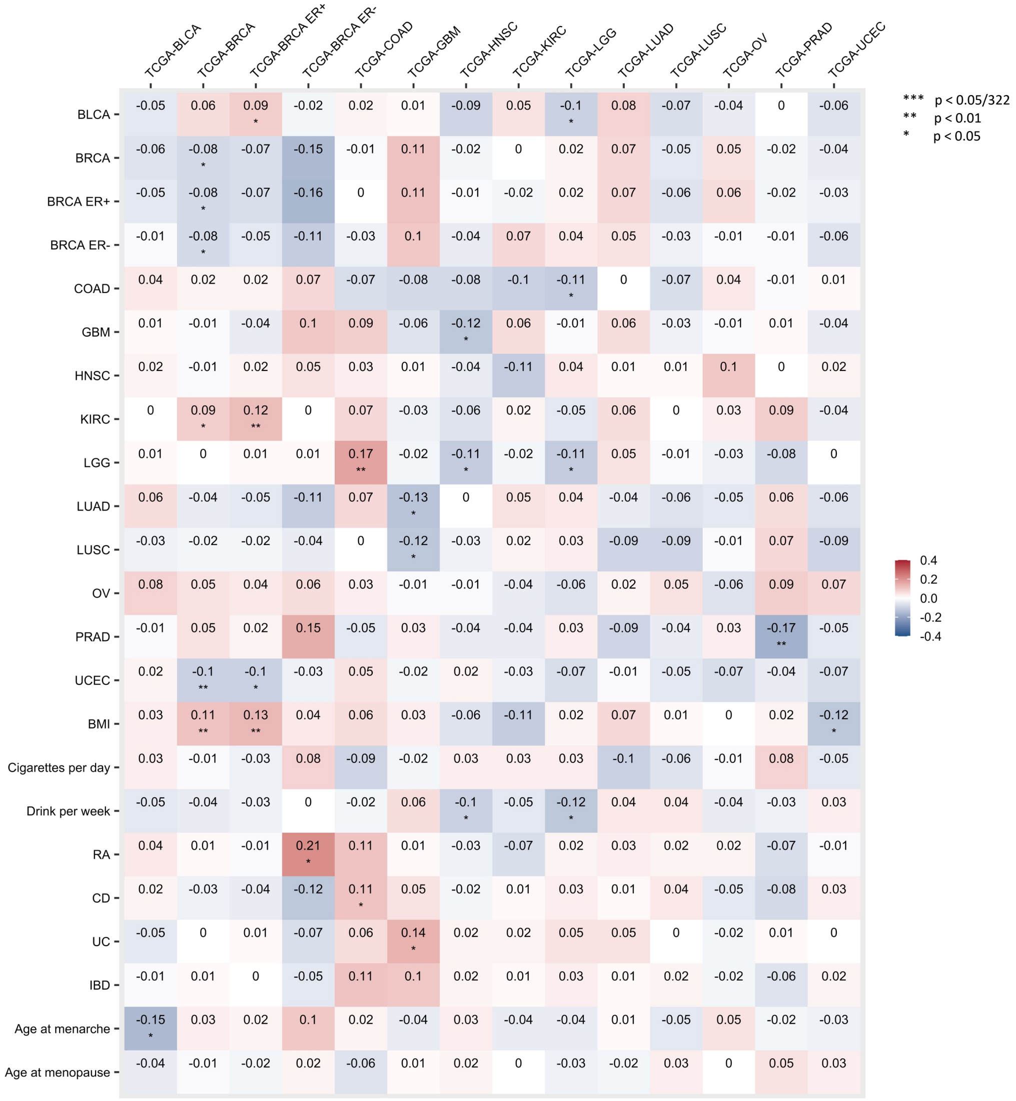


**Supplementary Fig. 1.** Correlations between germline PRS and age at cancer diagnosis across cancers. Number in each cell and the cell color represent the Spearman correlation (ρ) between germline PRS of cancers and non-cancer traits (y-axis) and age at cancer diagnosis in a cancer type (x-axis). Corrections passed the Bonferroni threshold ($p < 0.05/322 = 1.55 \times 10^{-4}$) are marked with triple asterisk (***), correlations with $p < 0.01$ are marked with double asterisk (**), and correlations with $p < 0.05$ are marked with single asterisk (*).

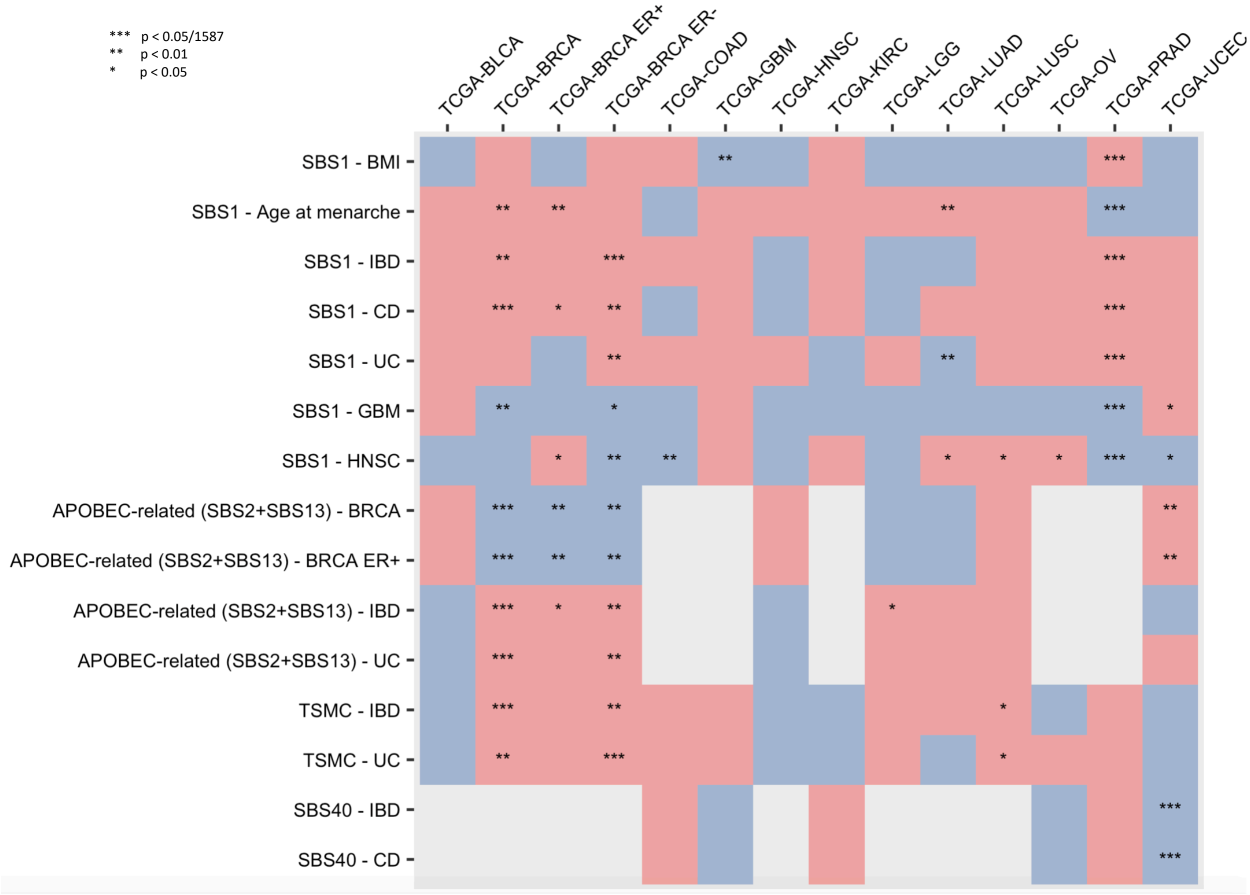


**Supplementary Fig. 2.** Significant associations between SBS signatures (or TSMC) and germline PRS across cancers. The cell color represents the direction of association between the number of somatic mutations of a SBS signature or TSMC and a germline PRS (y-axis) in a cancer type (x-axis): red = positive association; blue = inverse association. Associations passed the Bonferroni threshold ($p < 0.05/1587 = 3.15 \times 10^{-4}$) are marked with triple asterisk (***), associations with $p < 0.01$ are marked with double asterisk (**), and associations with $p < 0.05$ are marked with single asterisk (*).

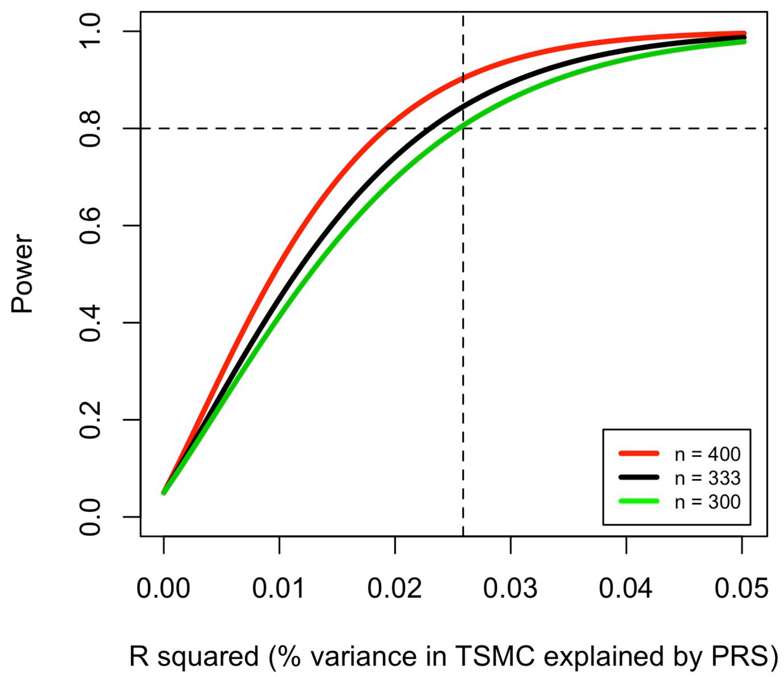


**Supplementary Fig. 3.** Power as a function of proportion of variance in TSMC explained by PRS for various sample sizes. Powers are calculated based on testing the association between PRS and TSMC for various proportions of variance and sample sizes at a type I error rate of 5%. The vertical dashed line represents the effect size for the rs2588809-TSMC association from Zhu et al[30]. We have at least 80% power (horizontal dashed line) to detect an association at (or greater than) the previously reported magnitude with sample size greater than 300.
